# Morphological differences between phyllosoma larvae from the western–central and eastern Pacific populations of the pronghorn spiny lobster *Panulirus penicillatus*

**DOI:** 10.1101/667469

**Authors:** Hirokazu Matsuda, Mitsuo Sakai, Takashi Yanagimoto, Seinen Chow

## Abstract

The pronghorn spiny lobster *Panulirus penicillatus* is known to have the widest distribution among palinurid lobster species, occurring in tropical and sub-tropical areas of the Indo-Pacific Ocean. In the Pacific Ocean, mitochondrial DNA analyses have revealed that the western–central and eastern populations are genetically isolated. We performed morphological investigations on mid- to late-stage phyllosoma larvae collected in these two areas. The larvae of the western–central population had a significantly narrower cephalic shield, shorter abdomen, and longer eyestalk than those of the eastern population. Additionally, for larvae larger than about 25-mm body length, the widest position of the cephalic shield in the western–central population was located closer to the middle of the median line of the cephalic shield than that in the eastern population. The ratio of width to length of cephalic shield and the ratio of cephalic shield width to thorax width are key traits for distinguishing between the larvae of the two populations.

## Introduction

The pronghorn spiny lobster *Panulirus penicillatus* occurs in tropical and sub-tropical areas of the Indo-Pacific Ocean. Its distribution is the widest among all species of spiny lobster, as there is no other spiny lobster species distributed across the Pacific Ocean (Holthuis 1991). The pronghorn spiny lobster is found in shallow water, usually at depths of less than 4 m, on rocky surfaces on the outer slopes of reefs and in water channels (Cockcroft *et al*. 2011). The pronghorn spiny lobster is an important target for coastal fisheries and supports the economy of many countries (Munro 2000). In many parts of its range, however, this species appears to be declining as a result of intense fishing pressure (Cockcroft *et al*. 2011). For the purpose of determining a sustainable catch for this lobster, it is necessary to perform adequate stock assessment and implement a fisheries management plan based on an understanding of the population structure.

Recently, mitochondrial DNA sequence analysis has been used to investigate the population structure of this lobster species in the Indo-Pacific Ocean (Chow *et al*. 2011, Abdullah *et al*. 2014a, b, Inacchei *et al*. 2016). *Panulirus penicillatus* in the Pacific Ocean has been divided into two distinct populations—the western–central (WC) population and the eastern (E) population—based on phylogenetic analyses of adult and phyllosoma larval samples collected throughout the Pacific Ocean (Chow *et al*. 2011, Abdullah *et al*. 2014a, Inacchei *et al*. 2016). Although this lobster has a long-lived teleplanic larval phase of at least 7–8 months, like many palinurid lobsters (Johnson 1968, Matsuda *et al*. 2006), the vast expanse of Pacific Ocean with no islands and no shallow substrate known as the East Pacific Barrier (EPB) appears to have effectively isolated these two populations. To date no notable differences have been found between the WC and the E populations in form or configuration of the body or appendages of adult lobsters, but individuals of the two populations have different body colors: greenish in the WC population and reddish-brown in the E population (George 2006, Abdullah *et al*. 2014a).

In the present study, we compared the body shape of *P. penicillatus* larvae collected in the western–central and eastern Pacific Ocean. This is the first report on regional differentiation of larval morphology in this spiny lobster species.

## Materials and Methods

### Collection of phyllosoma larvae

Phyllosoma samples for morphological examination were collected in three areas—the western Pacific (WP: station numbers 1–5 in Fig. 1 and Table 1), the central Pacific (CP: station 6), and the eastern Pacific (EP: station 7)—from the research vessels of the Fishery Agency of Japan and the Japan Fisheries Research and Education Agency from 2004 to 2016. Samples were collected by using an Isaacs-Kidd midwater trawl (IKMT: 8.7-m^2^ opening, 13 m long, 6-mm mesh, canvas cod-end) (Isaacs and Kidd 1953), Matsuda-Oozeki-Hu midwater trawl (MOHT: 5-m^2^ opening, 12 m long, 1.59-mm mesh, cod-end bucket) (Oozeki *et al*. 2004), a large midwater trawl (NST-660-SR) (MT: 161 m long, 10.4-mm to 60-mm mesh, 14.46-mm cod end) (Nichimo Co. Ltd., Tokyo, Japan), or a “larva capture” net (LC-100^2^-R3) (LC: 100-m^2^ opening, 36 m long, 6-mm mesh, cod-end bucket) (Nichimo Co. Ltd.) (Table 1). The towing velocity was 1–2.5 kn for the IKMT, 4–5 kn for the MOHT and MT, and 1–2 kn for the LC. Phyllosoma larvae sorted from the cod-end sample were preserved in 70% ethanol and transferred to the laboratory.

**Table 1.** Collection information for *Panulirus penicillatus* phyllosoma larvae used in this study.

| Station | Area of Pacific | Cruise No. | Period of cruise | Latitude (°N) | Longitude | Gear<sup>a</sup> | Depth range (m) | No. of larvae collected |
| --- | --- | --- | --- | --- | --- | --- | --- | --- |
| 1 | West | SHN04-5 | 6–12 Nov 2004 | 21°55'–27°30' | 124°07'–127°13' E | MOHT<sup>b</sup> | 0–300 | 34 |
| 2 | West | KY-08-04 | 28 Aug–2 Sep 2008 | 13°01'–14°11' | 142°03'–142°46' E | IKMT<sup>c</sup> | 0–300 | 16 |
| 3 | West | KY13-2 | 27 May–8 July 2013 | 11°00'–15°58' | 140°28'–143°30' E | IKMT<sup>c</sup> | 0–300 | 10 |
| 4 | West | KY14 | 26 Oct–27 Dec 2014 | 13°55'–28°05' | 124°51'–134°07' E | IKMT<sup>d</sup> | 0–200 | 10 |
| 5 | West | KY1504 | 22 Sep–31 Oct 2015 | 12°00'–18°00' | 128°59'–131°31' E | IKMT<sup>d</sup> | 0–200 | 10 |
| 6 | Central | KY155 | 25 Jan and 20 Feb 2016 | 26°35', 25°00' | 175°30' E, 159°59' W | MT<sup>e</sup> | 0–70 | 2 |
| 7 | East | KY07 | 1–2 Jan 2008 | 1°59', 3°59' | 95°59' W | LC<sup>f</sup> | 0–15 | 60 |

<sup>a</sup>MOHT, Matsuda-Oozeki-Hu midwater trawl; IKMT, Isaacs-Kidd midwater trawl; MT, large midwater trawl; LC, "larva capture" net
<sup>b</sup>Stepwise tows at about 300 m and 200 m during daytime and at about 80 m, 50 m and/or 25 m during nighttime. The net was kept at each depth for 5 to 15 minutes
<sup>c</sup>Oblique tows (20 cm/s) from around 300 m to the surface
<sup>d</sup>Oblique tows from around 200 m to the surface
<sup>e</sup>Horizontal tow during nighttime for 60 minutes
<sup>f</sup>Surface trawl (0–15 m) during nighttime for 30 minutes

**Fig. 1.**
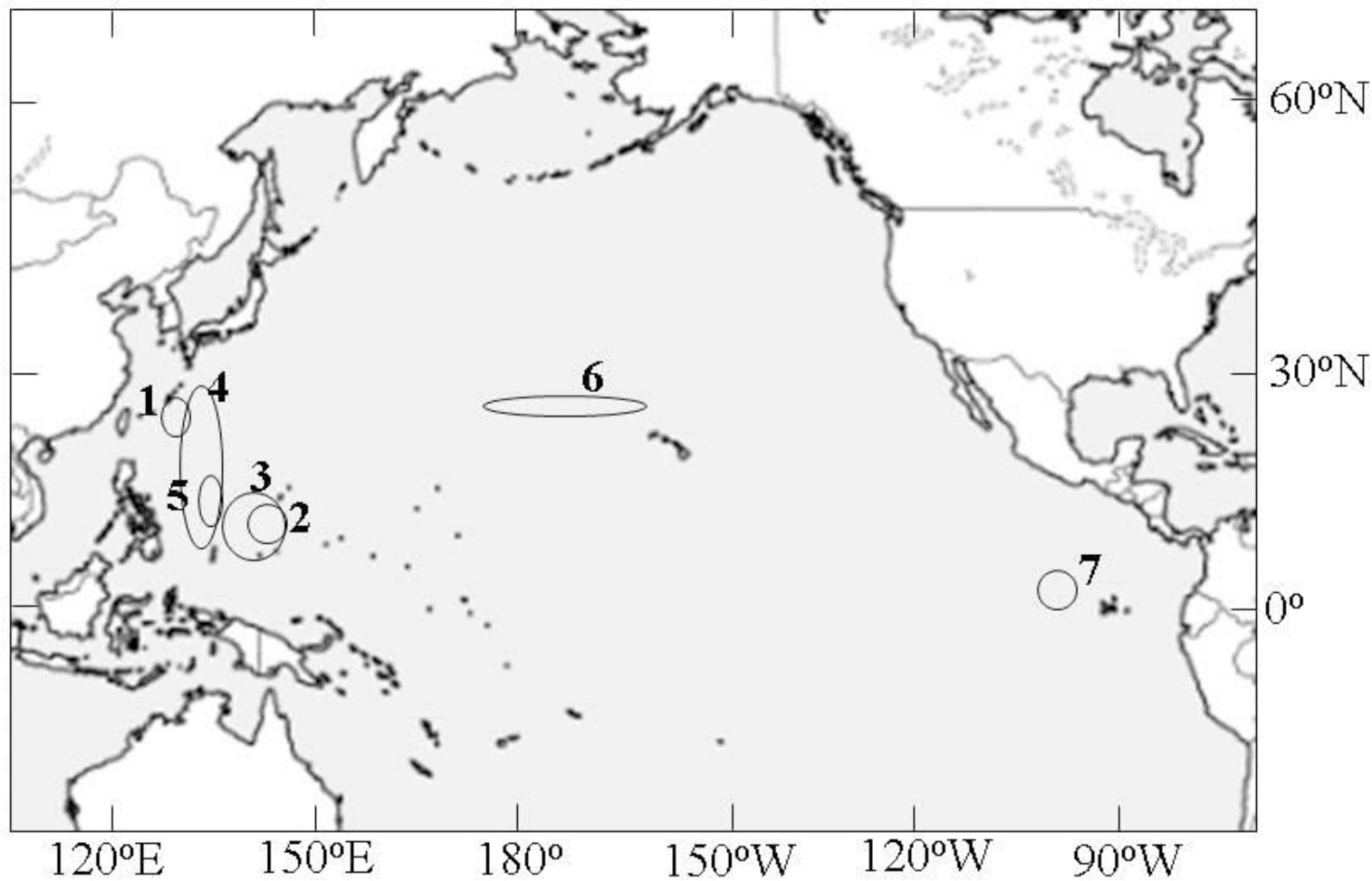
Collecting locations (open circles) for the phyllosoma larvae of *Panulirus penicillatus* in the Pacific Ocean. See Table 1 for detailed information.

### Identification of phyllosoma larvae

Phyllosoma larvae that were potentially *P. penicillatus* based on appearance were sorted out from the preserved samples using the morphological features given by Chow *et al*. (2006a) and Matsuda *et al*. (2006). The larval specimens selected on the basis of morphology were subjected to genetic analysis by nucleotide sequence or restriction fragment length polymorphism (RFLP) as previously described by Chow *et al*. (2006a, b). Larvae determined to be *P. penicillatus* from the DNA analysis were used for subsequent examination.

### Phyllosoma measurements and habitus analysis

For examinations of body habitus, larvae of the two *P. penicillatus* populations were measured using a Nikon profile projector (model V-12A, Nikon Ltd., Tokyo, Japan) as follows: body length (BL), from the anterior margin of the cephalic shield between the eyestalks to the posterior end of the abdomen; cephalic shield length (CL), from the anterior margin between the eyestalks to the posterior margin of the cephalic shield; cephalic shield width (CW), at the widest section of the cephalic shield; cephalic shield widest position (CWP), from the anterior margin of the cephalic shield to the point of intersection between the lines giving CL and CW; thorax width (TW), at the widest section of the thorax; abdominal length (AL), from a line level with the base of the abdomen to the posterior end of the abdomen; eyestalk length (ESL), from the anterior margin of the cephalic shield to the base of the eye (Fig. 2).

**Fig. 2.**
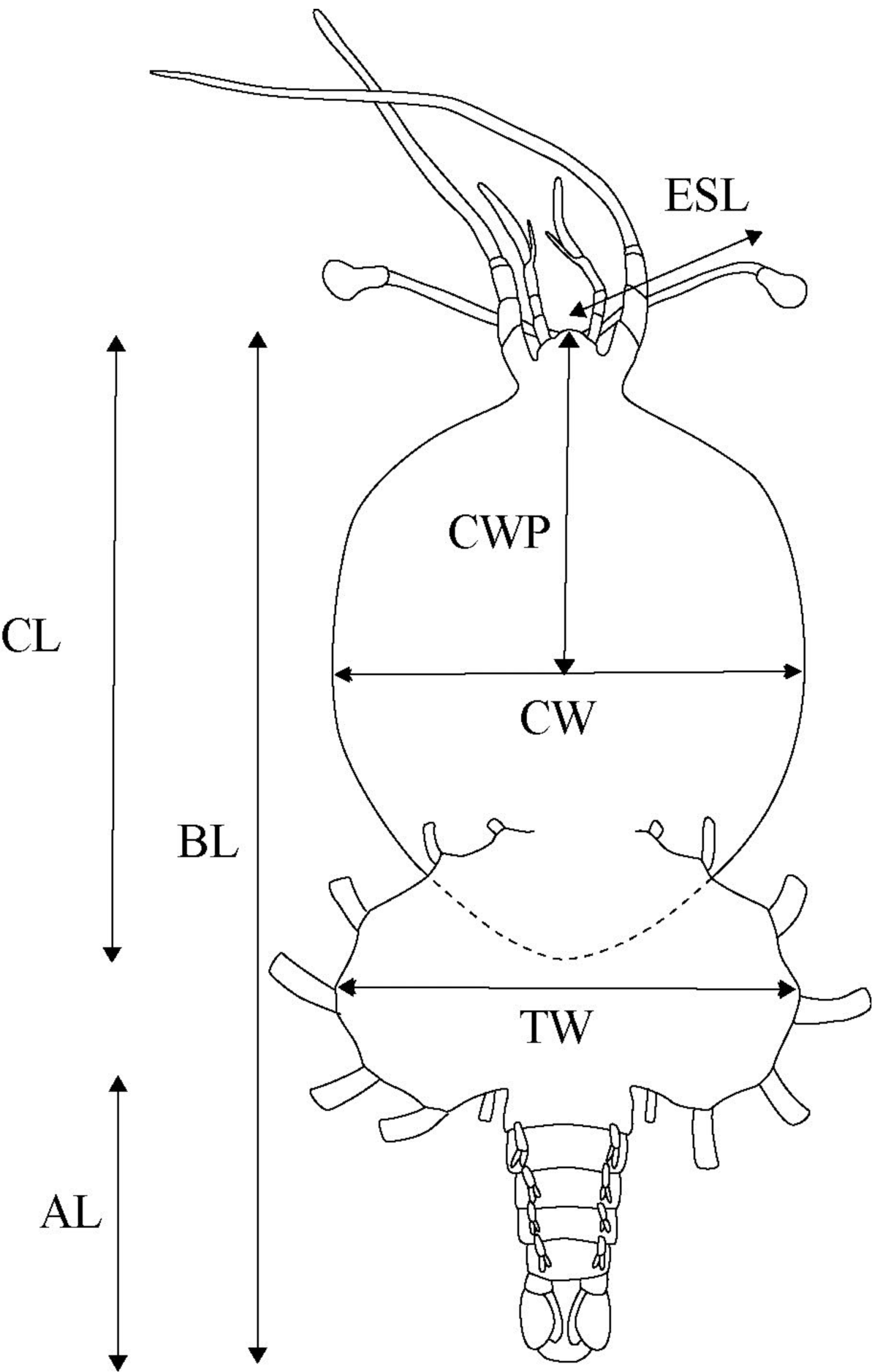
Diagram showing measurements of *Panulirus penicillatus* phyllosoma larvae. BL, body length; CL, cephalic shield length; CW, cephalic shield width; CWP, cephalic shield widest position; TW, thorax width; AL, abdomen length; ESL, eyestalk length.

To examine shifts in body shape with larval development, we analyzed allometric relationships between BL and body dimensions (CL, CW, TW, AL, and ESL) on the assumption that the relationships could be expressed with a logistic equation as follows:

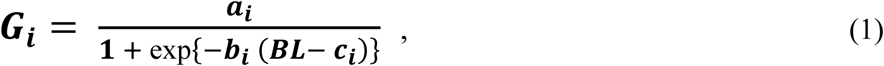

where *G*_*i*_ is the value of the *i*th larval body dimension (*i* = 1 for CL, *i* = 2 for CW, *i* = 3 for TW, *i* = 4 for AL, and *i* = 5 for ESL), and *a*_*i*_, *b*_*i*_, and *c*_*i*_ are parameters for the *i*th body dimension (to be estimated). A normal distribution 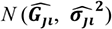 was assumed for describing the random error of each body dimension, where 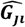 is the estimated *i*th body dimension for the *j*th datum of BL, and 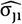 is the standard deviation. We assumed that ***σ***_***ji***_ is proportional to the following linear function of *BL*_*j*_:

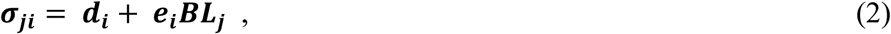

where *d*_*i*_ and *e*_*i*_ are parameters for adjusting the degree of the effect of *BL*_*j*_. The range from 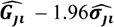 to 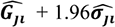 represents the estimated 95% confidence interval.

The parameters in equations (1) and (2) were estimated for each body dimension (*i* = 1 to 5) by maximizing the following log-likelihood function with the aid of non-linear optimization software:

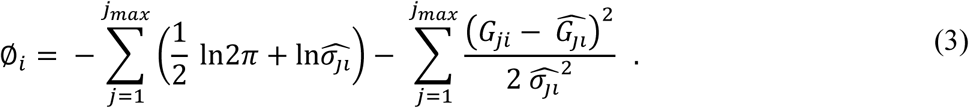

The relationships between BL and five body-dimension ratios (CW:CL, CW:TW, CWP:CL, AL:BL, and ESL:CL) representing the characteristics of body shape were assumed to be expressed by a first-order equation as follows:

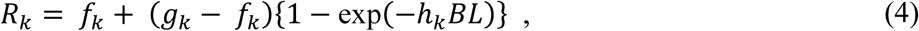

where *R*_*k*_ is the value of the *k*th body-dimension ratio (*k* = 1 for CW:CL, *k* = 2 for CW:TW, *k* = 3 for CWP:CL, *k* = 4 for AL:BL, and *k* = 5 for ESL:CL), and *f*_*k*_, *g*_*k*_, and *h*_*k*_ are parameters for the *k*th body-dimension ratio (to be estimated). A normal distribution 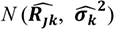 was assumed for describing the random error of each ratio, where 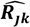 is the estimated *k*th body-dimension ratio for the *j*th datum of BL, and 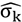 is the standard deviation. We assumed that 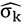 is constant and independent of *BL*_*j*_. The range from 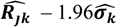 to 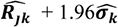 represents the estimated 95% confidence interval.

The parameters in equation (4) and *σ*_*k*_ were estimated for each body-dimension ratio (*k* = 1 to 5) by maximizing the following log-likelihood function with the aid of non-linear optimization software:

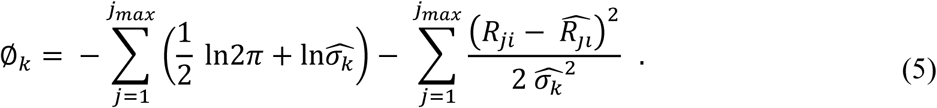

### Developmental stages

The larvae of the two populations were separated into developmental stages on the basis of staging criteria for cultured larvae of *P. penicillatus* (Matsuda *et al*. 2006). The body dimensions (BL, CL, CW, TW, AL, and ESL) for the larvae of the two populations were averaged for each morphological stage. The body-dimension ratios (CW:CL, CW:TW, CWP:CL, AL:BL, and ESL:CL) for the larvae from the WC and E populations were compared using a Mann–Whitney U test. The test was considered significant at *P* < 0.05.

## Results

The numbers of larvae determined to be *P. penicillatus* from morphologic features and subsequent genetic analysis were 80 from the WP, 2 from the CP, and 60 from the EP areas. Those from the WP and CP combined (*n* = 82) were determined to belong to the WC population, and the EP specimens (*n* = 60) were from the E population.

### Changes in body shape with development

The values for CL, CW, TW, AL, and ESL increased in a slight sigmoid curve with development (Fig. 3). Estimated parameters of the logistic equation for each body dimension are given in Table 2. The fitted lines that represent changes in the mean values of body dimensions as a function of BL show similar trends for CW and TW of the WC and E population larvae. For larvae larger than around 25-mm BL, on the other hand, CL and ESL of the WC population were shorter than in the E population, and AL of the WC population was greater than that of the E population.

**Table 2.** Estimated parameter values for the logistic equation 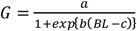 and the standard deviation *σ* (*σ* = *d* + *eBL*) for the allometric relationships between body length (*BL*) and body dimensions (*G*) for the phyllosoma larvae of the western–central(WC) and eastern (E) populations of *Panulirus penicillatus* in the Pacific Ocean. See text for explanation of each parameter.

| Parameter | Body dimension / Population |  |  |  |  |  |  |  |  |  |
| --- | --- | --- | --- | --- | --- | --- | --- | --- | --- | --- |
|  | CL |  | CW |  | TW |  | AL |  | ESL |  |
|  | WC | E | WC | E | WC | E | WC | E | WC | E |
| $a$ | 2.61 | 24.4 | 18.8 | 18.4 | 18.9 | 18.5 | 16.1 | 12.8 | 8.5 | 7.5 |
| $b$ | 0.112 | 0.109 | 0.125 | 0.134 | 0.118 | 0.111 | 0.132 | 0.163 | 0.124 | 0.118 |
| $c$ | 17.4 | 16.2 | 18.0 | 16.2 | 17.4 | 16.9 | 36.4 | 30.2 | 16.7 | 15.5 |
| $d$ | 0.127 | 0.000 | 0.134 | 0.020 | 0.000 | 0.000 | 0.000 | 0.000 | 0.000 | 0.001 |
| $e$ | 0.0119 | 0.0076 | 0.0110 | 0.0086 | 0.0201 | 0.0090 | 0.0116 | 0.0043 | 0.0159 | 0.0115 |

CL, cephalic shield length; CW, cephalic shield width; TW, thorax width; AL, abdomen length; ESL, eyestalk length

**Fig. 3.**
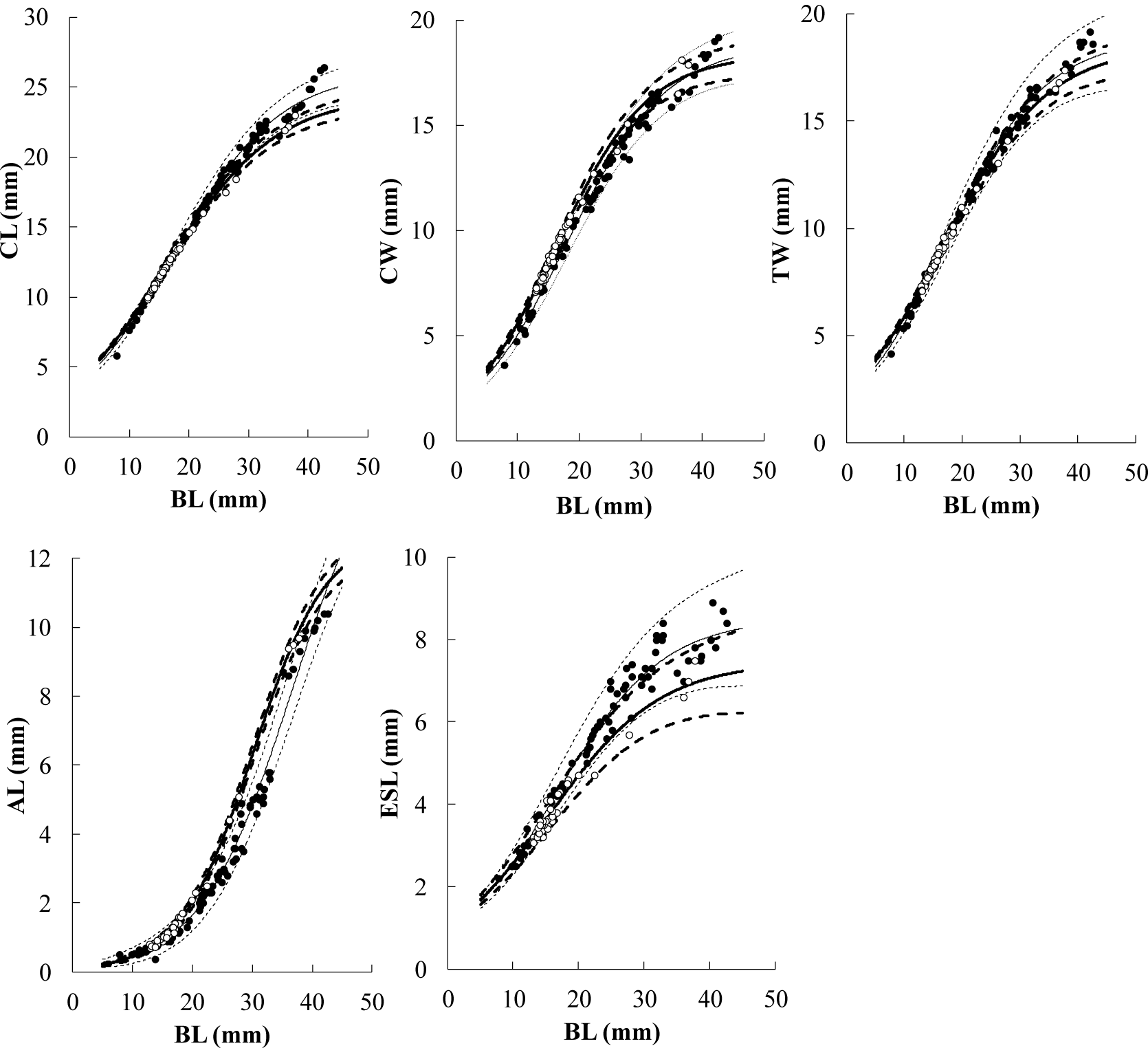
Allometric relationships between body length (BL) and body dimensions (CL, cephalic shield length; CW, cephalic shield width; TW, thorax width; AL, abdomen length; ESL, eyestalk length) in phyllosoma larvae of the two populations of *Panulirus penicillatus* in the Pacific Ocean. The thin and thick solid lines show the estimated mean values for the larvae in the western–central population (WC) and the eastern population (E), respectively. The 95% confidence intervals for respective mean values are shown by thin and thick dotted lines. Solid circles represent data for the WC population and open circles represent data for the E population.

The differences in body shape between larvae of the two populations were more conspicuous when body-dimension ratios (CW:CL, CW:TW, CWP:CL, AL:BL, and ESL:CL) were compared (Fig. 4). Estimated parameters of the first-order equation for each body-dimension ratio are given in Table 3. The values of CW:CL and CW:TW for the WC population larvae were lower than those for the E population larvae throughout the entire range of BL observed in the present study; there was barely any overlap in the 95% confidence intervals of CW:CL for the two populations for larvae larger than around 15-mm BL. The mean values of CW:CL and CW:TW for the WC population larvae over 30-mm BL were estimated as 0.73 and 1.00, respectively, and those in the E population were 0.79 and 1.06. CWP:CL for larvae larger than around 25-mm BL tended to be higher in the WC population than in the E population, with values of around 0.49 and 0.45, respectively. This indicates that the position of the widest part of the cephalic shield in the WC population larvae was close to the mid-point of the median line of the cephalic shield and that in the E population larvae it was located more anteriorly. The AL:BL values of the WC population larvae were lower than those of the E population larvae for larvae larger than around 20-mm BL. There was a trend of higher ESL:CL values in the WC population larvae than in the E population larvae, although there is considerable overlap in the 95% confidence intervals of the two populations.

**Table 3.** Estimated parameter values for the first-order equation *R* = *f* + (*g* − *f*) {1 − exp(−*hBL*)} and the standard deviation *σ* for the relationships between body length (BL) and body-dimension ratios (R) for the phyllosoma larvae of the western–central(WC) and eastern (E) populations of *Panuliruspenicillatus* in the Pacific Ocean. See text for an explanation of each parameter.

| Parameter | Body-dimension ratio / Population |  |  |  |  |  |  |  |  |  |
| --- | --- | --- | --- | --- | --- | --- | --- | --- | --- | --- |
|  | CW:CL |  | CW:TW |  | CWP:CL |  | ESL:CL |  | AL:BL |  |
|  | WC | E | WC | E | WC | E | WC | E | WC | E |
| $f$ | 0.376 | 0.000 | 0.572 | 0.000 | 1.088 | 0.552 | 0.341 | 0.009 | 0.006 | -0.030 |
| $g$ | 0.74 | 0.80 | 1.01 | 1.06 | 0.18 | 0.39 | 0.00 | 0.32 | -0.08 | -0.035 |
| $h$ | 0.120 | 0.179 | 0.133 | 0.218 | 0.253 | 0.028 | 0.000 | 0.325 | -0.033 | -0.017 |
| $\sigma$ | 0.0197 | 0.0145 | 0.0286 | 0.0201 | 0.0210 | 0.0136 | 0.0227 | 0.0155 | 0.0138 | 0.0055 |

CL, cephalic shield length; CW, cephalic shield width; TW, thorax width; CWP, cephalic shield widest position; ESL, eyestalk length; AL, abdomen length

**Fig. 4.**
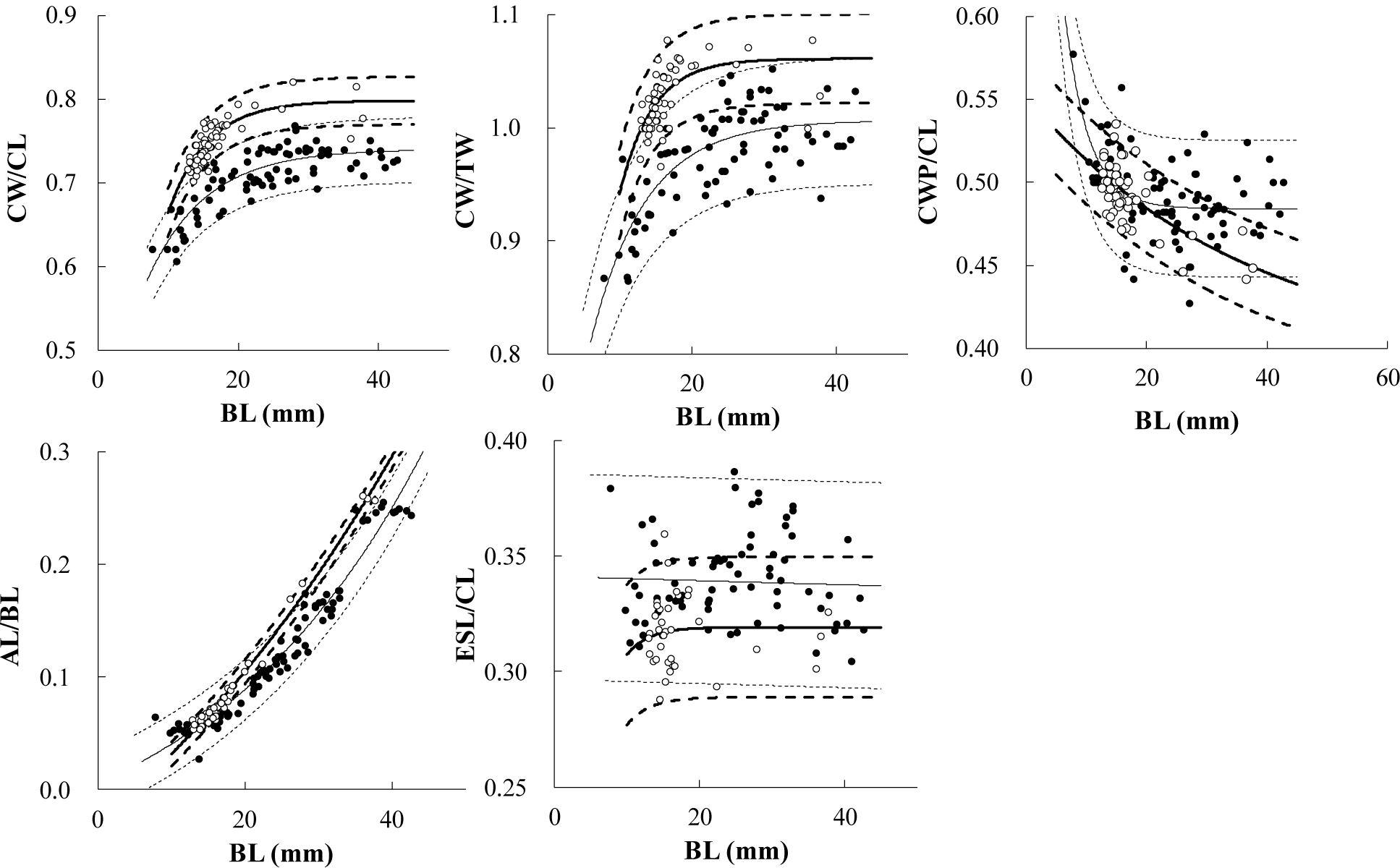
Relationships between body length (BL) and five body-dimension ratios (CW:CL, CW:TW, CWP:CL, AL:BL, and ESL:CL) in the phyllosoma larvae of two populations of *Panulirus penicillatus* in the Pacific Ocean. Thin and thick solid lines show estimated mean values for larvae of the western–central (WC) and eastern (E) populations, respectively. The 95% confidence intervals for the respective mean values are shown by thin and thick dotted lines. Solid circles represent data for the WC population and open circles represent data for the E population. CW, cephalic shield width; CL, cephalic shield length; TW, thorax width; CWP, cephalic shield widest position; AL, abdomen length; ESL, eyestalk length.

### Comparison of body dimensions of phyllosoma larvae from the WC and E populations by developmental stage

The 82 larvae of the WC population were assigned into 5 stages, from Stage VI to Stage X, and the 60 larvae of the E population were assigned into 4 stages, from Stage VII to Stage X. The various body dimensions for each stage are presented in Table 4. There were overlaps between the WC population and the E population in the range of each body dimension at all stages, with a tendency for BL, CL, and ESL to be higher in WC population larvae than in E population larvae.

**Table 4.** Comparison of morphometric sizes in stage VI to X phyllosoma larvae of the western–central (WC) and eastern (E) populations of *Panulirus penicillatus* in the Pacific Ocean. BL, body length; CL, cephalic shield length; CW, cephalic shield width; TW, thorax width; AL, abdomen length; ESL, eyestalk length.

|  |  | Stage / Population (No. of inds.) |  |  |  |  |  |  |  |  |  |
| --- | --- | --- | --- | --- | --- | --- | --- | --- | --- | --- | --- |
|  |  | VI |  | VII |  | VIII |  | IX |  | X |  |
|  |  | WC (10) | E (0) | WC (14) | E (48) | WC (25) | E (7) | WC (22) | E (2) | WC (11) | E (3) |
| BL | Mean | 11.1 | - | 15.3 | 15.1 | 23.0 | 19.3 | 29.6 | 26.9 | 39.0 | 36.8 |
|  | Min. | 7.8 | - | 11.1 | 12.8 | 17.5 | 17.7 | 24.2 | 26.1 | 35.0 | 36.0 |
|  | Max. | 12.4 | - | 17.8 | 17.7 | 28.1 | 22.4 | 32.8 | 27.8 | 42.6 | 37.7 |
| CL | Mean | 8.5 | - | 11.8 | 11.5 | 16.9 | 14.2 | 20.6 | 17.9 | 24.2 | 22.4 |
|  | Min. | 5.8 | - | 8.4 | 9.9 | 13.4 | 13.3 | 17.7 | 17.5 | 21.5 | 21.9 |
|  | Max. | 9.7 | - | 13.8 | 13.5 | 19.6 | 16.0 | 22.6 | 18.4 | 26.4 | 23.0 |
| CW | Mean | 5.5 | - | 8.0 | 8.5 | 12.2 | 11.0 | 15.1 | 14.4 | 17.6 | 17.5 |
|  | Min. | 3.6 | - | 5.1 | 7.1 | 9.3 | 10.2 | 13.1 | 13.8 | 15.9 | 16.5 |
|  | Max. | 6.1 | - | 9.5 | 10.2 | 15.0 | 12.7 | 16.6 | 15.1 | 19.2 | 18.1 |
| TW | Mean | 6.0 | - | 8.5 | 8.3 | 12.3 | 10.4 | 15.1 | 13.6 | 17.8 | 16.9 |
|  | Min. | 4.2 | - | 5.9 | 7.1 | 9.7 | 9.7 | 13.0 | 13.1 | 16.4 | 16.5 |
|  | Max. | 6.8 | - | 9.8 | 9.7 | 14.6 | 11.9 | 16.6 | 14.1 | 19.2 | 17.4 |
| AL | Mean | 0.6 | - | 0.9 | 1.0 | 2.4 | 1.9 | 4.6 | 4.8 | 9.6 | 9.5 |
|  | Min. | 0.5 | - | 0.4 | 0.7 | 1.2 | 1.6 | 2.8 | 4.4 | 8.6 | 9.4 |
|  | Max. | 0.7 | - | 1.2 | 2.3 | 3.6 | 2.5 | 5.8 | 5.1 | 10.4 | 9.7 |
| ESL | Mean | 2.8 | - | 3.8 | 3.6 | 5.8 | 4.6 | 7.1 | 5.7 | 7.9 | 7.0 |
|  | Min. | 2.2 | - | 1.9 | 3.1 | 4.4 | 4.5 | 4.5 | 5.7 | 7.0 | 6.6 |
|  | Max. | 3.4 | - | 4.5 | 4.3 | 7.4 | 4.7 | 8.4 | 5.7 | 8.9 | 7.5 |

WC population larvae had values of CW:CL and CW:TW that were significantly lower than E population larvae in almost all stages, indicating that the cephalic shield of the WC population larvae was relatively narrower than that of the E population larvae (Fig. 5, Table 5). The mean value for CWP:CL in Stage X of WC population larvae was 0.492 and significantly higher than that of E population larvae (0.453), also indicating that the widest position of cephalic shield of the WC population larvae was significantly closer to the central point of the median line of the cephalic shield than that in the E population larvae, which was located more anteriorly. There was a trend of shorter abdomens and longer eyestalks in the WC population larvae than in the E population larvae, as reflected by AL:BL and ESL:CL.

**Table 5.**
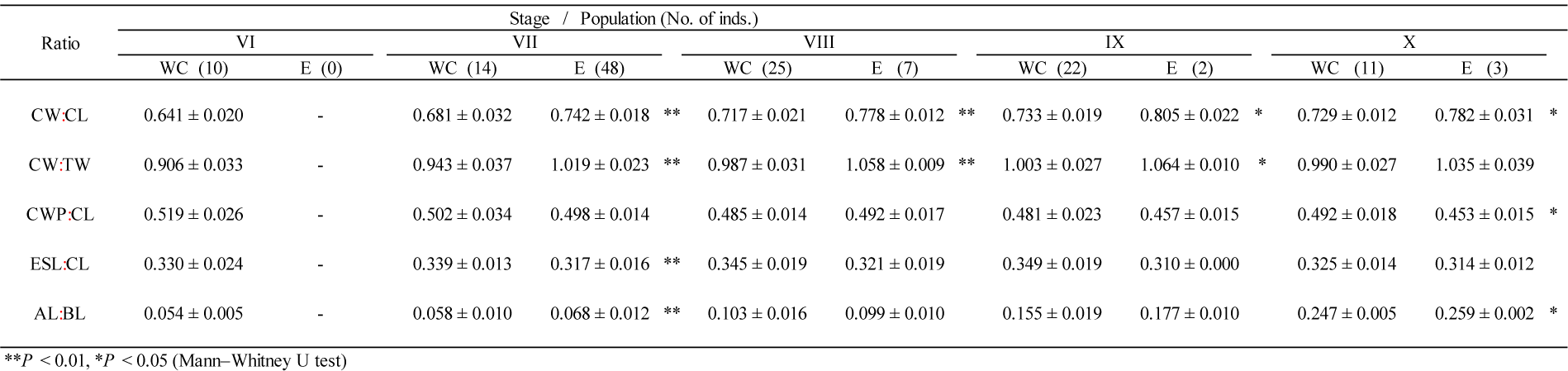
Comparison of body-dimension ratios in stage VI to X phyllosoma larvae of the western–central (WC) and eastern (E) populations of *Panulirus penicillatus* in the Pacific Ocean. BL, body length; CL, cephalic shield length; CW, cephalic shield width; CWP, cephalic shield widest position; TW, thorax width; AL, abdomen length; ESL, eyestalk length.

**Fig. 5.**
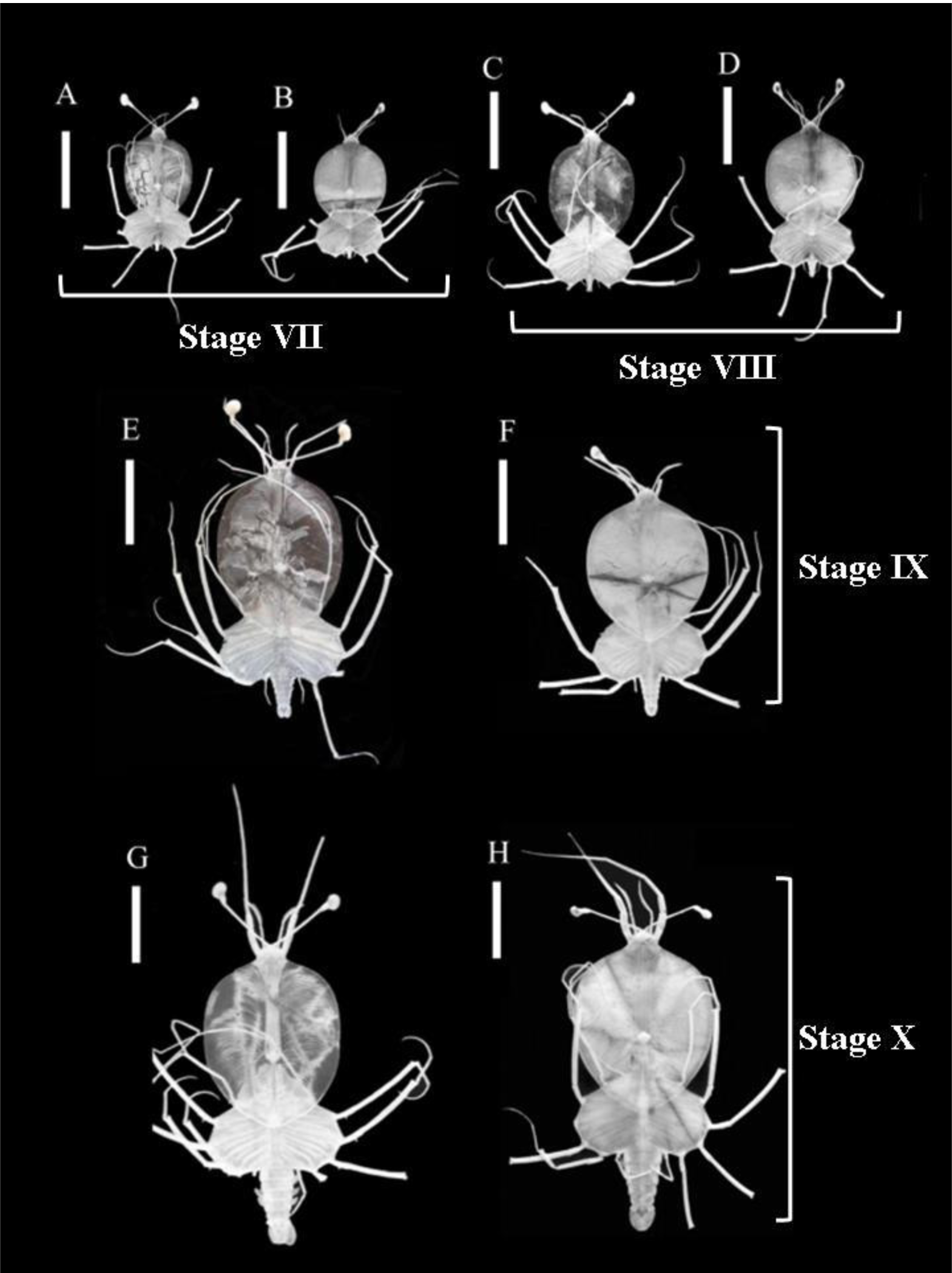
Phyllosoma larvae of *Panulirus penicillatus* from Stage VII to Stage X of the western–central population (A, C, E, and G) and the eastern population (B, D, F, and H) in the Pacific Ocean. Scale bar = 10 mm.

## Discussion

Our measurements of metric characteristics of *P. penicillatus* larvae show that phyllosoma larvae of the two populations in the Pacific Ocean have different body shapes: WC population larvae have a significantly narrower cephalic shield, shorter abdomen, and longer eyestalk than E population larvae. For larvae larger than 25-mm BL, the widest position of the cephalic shield in the WC population is closer to the central point of the median line of the cephalic shield than in the E population larvae, and that in the E population larvae was more anterior to the central point. Additionally, the ratios of cephalic shield width to cephalic shield length (CW:CL) and thorax width (CW:TW) were distinct between the larvae of the two populations: CW:CL and CW:TW of the WC population larvae were lower than those of the E population larvae over the entire BL range (10–40 mm) examined in this study.

The CW:TW ratio has been a key trait for identifying phyllosoma larvae of four species-groups of *Panulirus* lobsters (Groups I–IV), as proposed by George and Main (1967), and even species within the same group (McWilliam 1995, Chow *et al*. 2006a). For phyllosoma larvae larger than around 15-mm BL, those of *P. penicillatus* belonging to Group II had a body shape with CW:TW values of around 1.0, significantly higher than those of Group I lobsters and lower than those of Group IV lobsters (Chow *et al*. 2006a). The present study confirms that the CW:TW ratio is even applicable for distinguishing the two populations of *P. penicillatus* larvae. The CW:CL ratio was also valid for separating the larvae of the two *P. penicillatus* populations, because there was little overlap in the 95% confidence intervals of this ratio between the two populations in larvae larger than around 15-mm BL.

Polygenetic analysis of mtDNA has confirmed the prolonged historical isolation between the WC and E populations; the EPB—the 4000- to 7000-km expanse of deep water without islands—appears to have effectively isolated the two populations (Chow *et al*. 2011; Abdullah *et al*. 2014a; Inacchei *et al*. 2016). Adult lobsters of the two populations on each side of the EPB have different body colors: greenish in the WC population and reddish-brown in the E population (George 2006, Abdullah *et al*. 2014a). Genetic analysis suggests that the two populations remain as sub-species of *P. penicillatus* (Abdullah *et al*. 2014a). The discrepancies in the values of CW:TW between the larvae of the two populations (e.g. 1.003 [WC] vs. 1.064 [E] in the penultimate phyllosoma stage) are much larger than that between the larvae of subspecies in the *P. longipes* complex collected in the same area of the western Pacific Ocean (*P. l. longipes* [0.852] vs. *P. l. bispinosus* [0.852] at the same stage) (Chow *et al*. 2006a).

The morphological differences observed between the larvae of the two populations of *P. penicillatus* might be genetically determined characteristics. However, environmental factors can affect larval morphology. The shape of cephalic shield is known to differ between wild and laboratory-produced larvae. The final phyllosomal stage (Stage X) of the WC population of *P. penicillatus* cultured in the laboratory had a mean CW:TW value of 0.953 (Matsuda *et al*. 2006), whereas the values for wild-caught larvae of the WC population were 0.990 (this study), 1.00 (Chow *et al*. 2006a), and 1.01 (Johnson 1968). There was a similar discrepancy between wild and laboratory-produced larvae of *P. longipes bispinosus* and *P. japonicus*. The mean CW:TW values of cultured stages VII to X larvae of *P. l. bispinosus* were 0.784, 0.779, 0.820, and 0.813 (Matsuda and Yamakawa 2000), whereas those of wild larvae were 0.821, 0.838, 0.852, and 0.836 (Chow *et al*. 2006a). The mean CW:TW values of cultured stages VII to X larvae of *P. japonicus* were 0.688, 0.697, 0.710, and 0.749 (Matsuda 2005), whereas those of wild larvae were 0.740, 0.761, 0.761, and 0.766 (Chow *et al*. 2006a). Mid- to late-stage wild phyllosomae of *P. japonicus* also had a wider cephalic shield than cultured larvae (Hamasaki *et al*. 2012).

Unfortunately there are no studies into environmental factors that might affect the shape of the cephalic shield, although there must be discrepancies between the environments in a tank and in the wild (e.g. water current, temperature, light, and food). It will be necessary to culture phyllosoma larvae of the two populations under the same conditions to determine whether genetic or environmental factors induce the difference in the shape of the cephalic shield.

Several publications have described *P. penicillatus* phyllosoma larvae collected in the ocean. In the Pacific Ocean, Johnson (1968) caught 128 larvae of this species in Hawaiian and neighboring waters (central Pacific Ocean) and described some developmental stages. His later-stage larvae have CW:CL values of 0.75, indicating that the body shape of his specimens deviates from the typical one found in the present study for the WC population of *P. penicillatus*. In the present study, we only collected two *P. penicillatus* larvae from the waters near the Hawaiian archipelago; they were at Stage VII and Stage X. The CW:CL and CW:TW ratios of the Stage X larva were 0.751 and 1.035, which are near the upper limits of the 95% confidence intervals for the WC population larvae. As this discussion suggests, it may be difficult to distinguish between the larvae from Hawaiian and neighboring waters and those of the E population.

Johnson (1971a) reported a Stage VIII phyllosoma larva (21-mm BL) caught in the South China Sea. The body shape corresponds to the typical body shape of the WC population larvae in the present study. Moreover, a Stage X larva (32 mm), equivalent to our Stage IX, collected from the east side of the EPB off the coast of Mexico, was described and illustrated by Johnson (1971b). This larva had a body shape typical of E population larvae.

There are several reports of *P. penicillatus* phyllosoma larvae from the Indian Ocean. A phyllosoma larva (19.8-mm BL, our Stage VIII) caught off the Natal coast of South Africa had CW:CL and CW:TW ratios of 0.766 and 1.017, respectively, seemingly closer to the body shape of E population larvae than of WC population larvae (Berry 1974). Prasad *et al*. (1975) roughly illustrated a sequence of developmental phyllosoma stages from Stage VII to Stage XII from the tropical and southwestern areas of the Indian Ocean. Their Stage XII, corresponding to our Stage X, has CW:CL and CW:TW ratios of 0.746 and 0.961, respectively, appearing mostly like the WC population larvae. Tampi and George (1975) also show a sketch of a final-stage larva (35.5-mm BL) from the eastern Indian Ocean; the larva has CW:CL and CW:TW values of 0.732 and 1.050, respectively, and the body shape appears to have characteristics of both WC and E population larvae. Abdullah *et al*. (2014b) observed distinct genetic isolation of populations at the northwestern and southwestern edges of the distribution of *P. penicillatus* in the Indian Ocean, and gene flow within the population in the central and eastern regions of the ocean. Therefore, there is a probably a wide range of variation in the larval morphology of *P. penicillatus* in the Indo-Pacific Ocean.

## Acknowledgments

We are grateful to the captains and crew members of R/V *Shun-yo Maru* of the Japan Fisheries Research and Education Agency, and R/V *Kaiyo Maru* of the Fisheries Agency of Japan for supporting sample collection. This work was supported in part by a Grant-in-Aid for Scientific Research on Priority Areas (B) (No. 25450292) from the Ministry of Education, Science, Sports, and Culture of Japan.

## Literature Cited

Abdullah, M.F., Chow, S., Sakai, M., Cheng, J., & Imai, H., 2014a. Genetic diversity and population structure of pronghorn spiny lobster *Panulirus penicillatus* in the Pacific region. Pacific Science, 68:197–211.

Abdullah, M.F., Muththalib, M., Salama, A.J., & Imai, H., 2014b. Genetic isolation among the Northwestern, Southwestern and Central-Eastern Indian Ocean populations of the pronghorn spiny lobster *Panulirus penicillatus*. International Journal of Molecular Science, 15:9242–9254.

Berry, P.F., 1974. Palinurid and scyllarid lobster larvae of the Natal Coast, South Africa. Oceanographic Research Institute Investigational Report, 34: 1–44.

Chow, S., Yamada, H., & Suzuki, N., 2006a. Identification of mid-to final stage phyllosoma larvae of the genus *Panulirus* White, 1847 collected in the Ryukyu Archipelago. Crustaceana 79:745–764.

Chow, S., Suzuki, N., Imai, H., & Yoshimura, T. 2006b. Molecular species identification of spiny lobster phyllosoma larvae of the genus *Panulirus* from the Northwestern Pacific. Marine Biotechnology, 8:260–267.

Chow, S., Jeffs, A., Miyake, Y., Konishi, K., Okazaki, M., Suzuki, N., Abdullah, M.F., Imai, H., Wakabayashi, T., & Sakai, M., 2011. Genetic isolation between the western and eastern Pacific populations of pronghorn spiny lobster *Panulirus penicillatus*. PloS ONE, 6(12): e29280.

Cockcroft, A., MacDiarmid, A., & Butler, M., 2011. “*Panulirus penicillatus*”. The IUCN Red List of Threatened Species. International Union for Conservation of Nature and Natural Resources. (URL:http://dx.doi.org/10.2305/IUCN.UK.2011-1.RLTS.T169951A6691002.en.)

George, R.W., 2006. Tethys sea fragmentation and speciation of *Panulirus* spiny lobsters. Crustaceana, 78:1281–1309.

George, R.W., & Main, A.R., 1967. The evolution of spiny lobsters (Palinuridae): a study of evolution in the marine environment. Evolution, 21: 803–820.

Hamasaki, K., Mizumoto, Y., Jinbo, T., & Murakami, K., 2012. Ontogenetic change of body density and shape of the phyllosoma larvae of the Japanese spiny lobster *Panulirus japonicus*. Journal of Crustacean Biology, 32(3): 395–404.

Holthuis, L.B., 1991. Marine lobsters of the world. An annotated and illustrated catalogue of species of interest to fisheries known to date. FAO Fisheries Synopsis, 125(13): 151–152.

Inacchei, M., Gaither, M.R., Bowen, B.W., & Toonen, R.J., 2016. Testing dispersal limits in the sea: range-wide phylogeography of the pronghorn spiny lobster *Panulirus penicillatus*. Journal of Biogeography, 43: 1032–1044.

Isaacs, J.D., & Kidd, L.W., 1953. Isaacs-Kidd midwater trawl. Final Report. Scripps Institute of Oceanography. Reference 53-3, Oceanographic Equipment Report No. 1.

Johnson, M.W., 1968. Palinurid phyllosomas from the Hawaiian archipelago (Palinuridae). Crustaceana, Supplement 2: 59–79.

Johnson, M.W., 1971a. On palinurid and scyllarid lobster larvae and their distribution in the South China Sea (Decapoda, Palinuridae). Crustaceana, 21: 247–282.

Johnson, M.W., 1971b. The palinurid and scyllarid lobster larvae of the tropical eastern Pacific and their distribution as related to the prevailing hydrography. Bulletin of the Scripps Institution of Oceanography, 19: 1–36.

Matsuda, H., & Yamakawa, T., 2000. The completedevelopment and morphological changes of larval *Panulirus lomgipes* (Decapoda, Palinuridae) under laboratory conditions. Fisheries Science, 66: 278–293.

Matsuda, H., 2005. Studies on the larval culture and development of *Panulirus* lobsters. PhD Thesis, Kyoto University, Kyoto, Japan, pp. 1–211.

Matsuda, H., Takenouchi, T., & Goldstein, J.S., 2006 The complete larval development of the Pronghorn spiny lobster *Panulirus penicillatus* (Decapoda: Palinuridae) in culture. Journal of Crustacean Biology, 26:579–600.

McWilliam, P.S., 1995. Evolution in the phyllosoma and puerulus phases of the spiny lobster genus *Panulirus* White. Journal of Crustacean Biology, 15 (3): 542–557.

Munro, J.L., 2000. Fisheries for spiny lobsters in the tropical Indo-West Pacific. In: B.F. Phillips & J. Kittaka, (eds.), Spiny Lobsters Fisheries and Culture. Second edition, Fishing News Books, Oxford, pp. 90–97.

Oozeki, Y., Hu, F.X., Kubota, H., Sugisaki, H., & Kimura, R., 2004. Newly designed quantitative frame trawl for sampling larval and juvenile pelagic fish. Fisheries Science, 70:223–232.

Prasad, R.R., Tampi, P.R.S., & George, M.J., 1975. Phyllosomas from the Indian Ocean collected by the Dana Expedition 1928-1930. Journal of the Marine Biological Association of India, 17: 56–107.

Tampi, P.R.S., & George, M.J., 1975. Phyllosoma larvae in the IIOE (1960 – 65) collections – systematics. Mahasagar, 8: 15–44.

